# Panx1b modulates the luminance response and direction of motion in the zebrafish

**DOI:** 10.1101/2021.07.21.453251

**Authors:** Nickie Safarian, Sarah Houshangi-Tabrizi, Christiane Zoidl, Georg R. Zoidl

## Abstract

Pannexin1 (Panx1) can form ATP-permeable integral membrane channels that play roles in the physiology of the visual system. Two independent gene copies of Panx1*, panx1a* and *panx1b,* have been identified in the zebrafish with unique and shared properties and tissue expression patterns. Panx1a channels, located in horizontal cells of the outer retina, modulate light decrement detection through an ATP/pH-dependent mechanisms and adenosine/dopamine signaling. Here, we decipher how the strategic localization of Panx1b channels in the inner retina and ganglion cell layer modulates visually evoked motor behavior. We describe a *panx1b* knockout model generated by TALEN technology. The RNA-seq analysis of 6 days post-fertilization larvae is confirmed by Real-Time PCR and paired with testing of visual-motor behaviors. The Panx1b protein emerges as a modulator of the circadian clock system. The loss of *panx1b* also disrupts the retinal response to the abrupt loss of illumination and decreases the larval ability to follow leftward direction of motion in the dark. The evidence suggests that in the retina Panx1b contributes to the OFF pathways function, like Panx1a, though through different signaling mechanisms. In this process, the loss of Panx1b channels compromises the final output of luminance as well as direction of motion detector RGCs. In addition, the disruption of the circadian clock system in mutants suggests that Panx1b could participate in non-image forming processes in the inner retina.

## Introduction

Pannexin 1 (Panx1) forms integral membrane channels generally assumed to have roles in the release of ATP. The cryoEM structure of Panx1 as a heptameric selective chloride channels was resolved recently (Deng, He et al. 2020, Jin, Zhang et al. 2020, Michalski, Syrjanen et al. 2020, Mou, Ke et al. 2020, Qu, Dong et al. 2020, Ruan, Orozco et al. 2020, Mim, Perkins et al. 2021). The significant expression of Panx1 in the mouse eye has fostered research on functional implications in the normal retina physiology and pathologies related to ocular hypertension or ischemia in mouse models (Ray, Zoidl et al. 2005, Dvoriantchikova, Ivanov et al. 2006, Dvoriantchikova, Ivanov et al. 2012, Kranz, Dorgau et al. 2013, Pronin, Pham et al. 2019). In parallel to ongoing work in rodents, the accessibility of the zebrafish at all stages of development to adulthood in combination with the possibility of behavioral assessment of visual system functions allows to study the mechanisms which involve Panx1 in vision using this complementary animal model.

The zebrafish has two copies of *panx1*, *panx1a* and *panx1b*, which resulted from the teleost third whole-genome duplication (R3 WGD) event occurring between 320 and 350 million years ago (MYA) (Bond, Wang et al. 2012, Ravi and Venkatesh 2018). The two Panx1 genes show distinct tissue distribution patterns and have shared and distinct biophysical properties (Prochnow, Hoffmann et al. 2009, Bond, Wang et al. 2012, Kurtenbach, Prochnow et al. 2013). The *panx1b* copy shows higher expression levels in the brain and eyes, compared to *panx1a* (Bond, Wang et al. 2012, Kurtenbach, Prochnow et al. 2013). In the zebrafish retina, Panx1b’s immunoreactivity is prominent in the inner retina and ganglion cells (GCs) (Kurtenbach, Prochnow et al. 2013), whereas Panx1a is localized on the surface of horizontal cell dendrites invaginating cone pedicles (Prochnow, Hoffmann et al. 2009). Other prominent biophysical differences are the smaller unitary conductance of Panx1b channels and longer activation time when compared to Panx1a (Kurtenbach, Prochnow et al. 2013).

To the best of present knowledge, the differences suggest a functional specialization in neuronal circuits shaping visual output of the retina. First evidence was found in the outer retina, where Panx1 channels affects a slow ATP/pH-mediated mechanism and Cx55.5 a fast ephaptic mechanism tuning feedback from horizontal cells to cones (Cenedese, de Graaff et al. 2017). Next, we showed that Panx1a modulates the detection of light decrements through the retinal OFF-pathway using adenosine and dopaminergic signalling pathways (Safarian, Whyte-Fagundes et al. 2020). Here, the roles of Panx1b channels in the inner retina were addressed using a *panx1b* knockout model generated with engineered transcription activator-like effector nucleases (TALEN) (Bedell, Wang et al. 2012). RNA-seq data of 6 days post-fertilization (dpf) larvae were confirmed by Real-Time PCR and paired with testing of visual-motor behaviors. The results showed that the loss of Panx1b disrupts the retinal response to the abrupt loss of illumination and causes a decreased visual acuity for detecting leftward motion in the dark. This outcome suggests a role of Panx1b in retinal OFF-pathways and directional coding retinal ganglion cell populations. The Gene Ontology (GO) enrichment analysis showed that different to Panx1a, Panx1b does not affect the adenosine-dopamine signaling pathway. Instead, Panx1b emerged as a modulator of the circadian clock system. Since roles of Panx1 in the sleep/wake cycle of rodents have been described we propose similar functions of Panx1b in non-image forming processes of the zebrafish visual system.

## Results

### Targeted disruption of the panx1b gene

For TALEN-mediated genome editing, a single HindIII restriction site in the 4th coding exon of *panx1b* was targeted (Fig. 1a). Twenty-five picograms (pg) of a pair of TALEN cRNAs was microinjected into one-cell stage embryos. A restriction fragment length polymorphism (RFLP) test of 17 randomly selected larvae at six-day post fertilization (6dpf) revealed a gene editing efficiency of ≈ 88%, as evidenced by a partial loss of the Hind III restriction enzyme recognition sequence located in the spacer region between the TALEN binding sites (Fig. 1b). The DNA sequence analysis of multiple F0 larvae confirmed the efficient generation of deletions of varying lengths (Fig. 1c). A positive adult F0 founder transmitting an 11-base pair (bp) frameshifting deletion (*panx1b^Δ11^*) was selected to establish the knockout line (*panx1b−/−*) used for further experimentation. The eleven bp deletion caused a frameshift at amino acid E179 of the Panx1b protein, resulting in a premature stop codon. The truncated 179-amino-acid protein *Panx1b^Δ11^* lacks the transmembrane regions 3 and 4, and the entire carboxyterminal domain. The lack of these critical domains suggested that the mutant was not capable of forming functional Panx1 channels (Fig. 1d).

**Figure 1.**
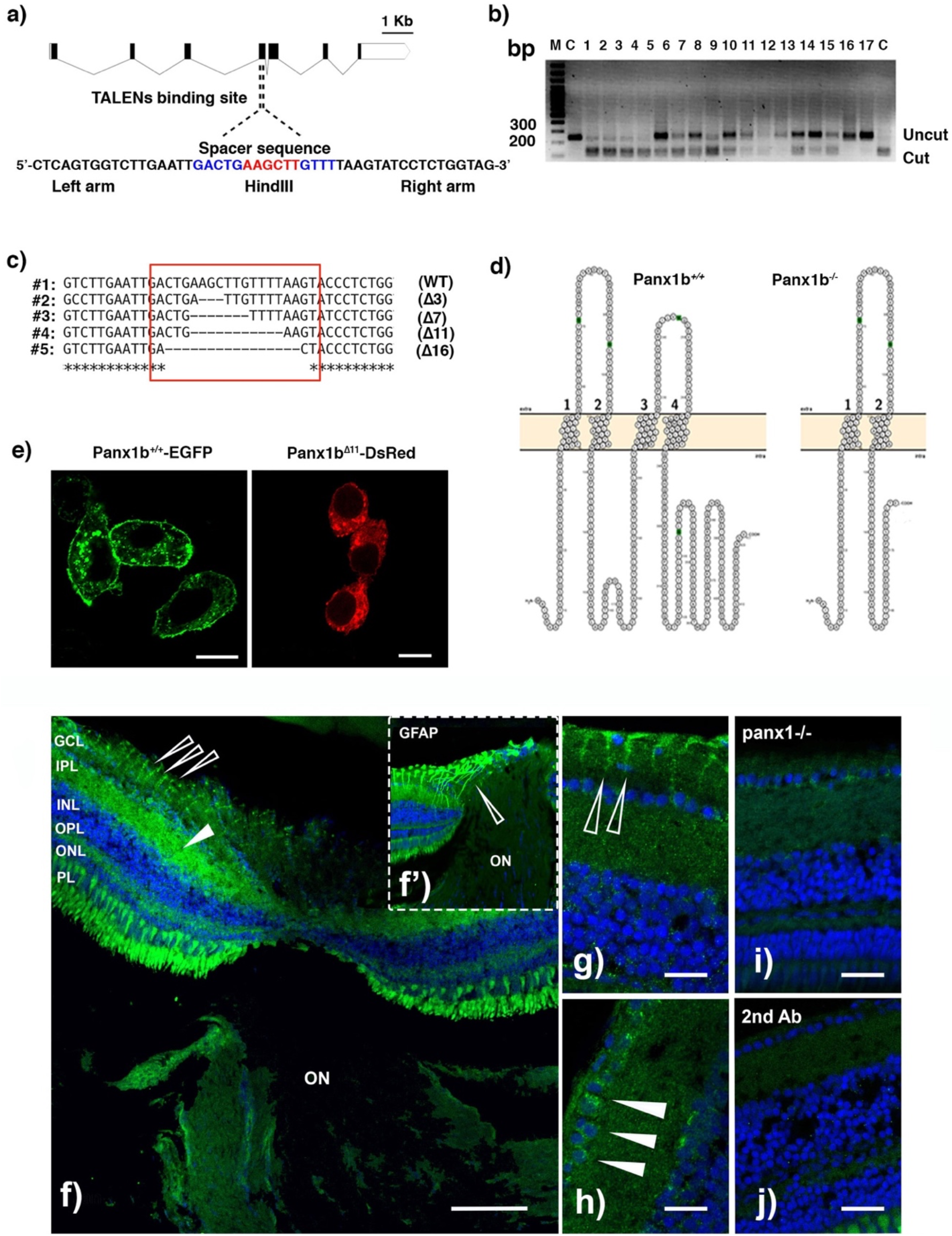
Generating panx1b^−/−^ fish using TALENs. **a)** Cartoon depicting the zebrafish *panx1b* gene structure with seven coding exons. The left and right TALENs binding sequence with the spacer sequence and HindIII restriction site in blue and red is highlighted. **b)** The RFLP-assay shows the loss of the HindIII recognition sequence (uncut) in fourteen out of seventeen F0 larvae tested. **c)** The sequence alignment demonstrates small (3 to 16 bp) deletions causing frameshift mutations (in red box) in four different F0 larvae. **d)** The Protter open-source tool *(wlab.ethz.ch/protter)* visualized the predicted topology of the wild type and mutant *Panx1b* proteins. The 11 bp deletion in *Panx1b* exon 4 resulted in a frameshift causing a premature stop codon at amino acid E179. **e)** Localization of the truncated *Panx1b* protein – Confocal images of transiently transfected *Panx1b^+/+^-EGFP* (left panel) and *Panx1b^Δ11^-DsRed-monomer* (right panel) proteins in Neuro2a cells. Scale bar: 10 μm. **f)** *Panx1b* immunoreactivity in the adult retina of TL fish. The section represents the retina at the contact point with the optic nerve (ON). The closed triangle points at Panx1b staining in the IPL. The open triangle represents Panx1b immunoreactivity in end-feet of Muller glia. The inset f’) shows GFAP-stained Muller glia in the same region indicated by an open triangle. **g)** Open arrows point at Panx1b-positive Muller glia end feet. **h)** Closed triangles point at Panx1b positive ganglion cells. **i)** Lack of significant Panx1b immunoreactivity in the adult *panx1b^−/−^* retina. **j)** control with 2^nd^ antibody only. Please note that the gain in green channel in images **i)** & **j)** was enhanced to visualize background staining. Abbreviations: WT, wild type TL; GCL, ganglion cell layer; IPL, inner plexiform layer; INL, inner nuclear layer; OPL, outer plexiform layer; ON, optic nerve. Scale bars: f = 100μm; g & h = 25μm; I & J = 20μm.

After transient transfection into mouse Neuroblastoma 2a (Neuro2a) cells, the *Panx1bΔ-DsRed-monomer* protein was detected in the cytoplasm. No significant signal in the cell membrane was detected, implying that the mutant protein was most likely unable to traffic efficiently to the cell membrane (**Fig. 1e**, left panel). The *Panx1b^wt^-EGFP* protein was detectable in the perinuclear region and the plasma membrane (**Fig. 1e**, right panel), in line with previous reports (Prochnow, Hoffmann et al. 2009, Prochnow, Hoffmann et al. 2009, Kurtenbach, Prochnow et al. 2013). The Panx1b immunoreactivity in the adult zebrafish retina was found mainly in the inner plexiform layer, and in structures resembling the end feet of Muller glia, or ganglion cells (**Fig. 1f-h**), corroborating the previously demonstrated localization identified with a different antibody (Kurtenbach, Prochnow et al. 2013). The Panx1b immunoreactivity was reduced in *panx1b^−/−^*, and absent when the primary antibody was omitted during the staining procedure **(Fig.1i,j)**.

### Characterization of panx1b^−/−^ larvae

The *panx1b^−/−^* survival rate was significantly lower than TL controls at 1 dpf (values in %: TL, 80; *panx1b^−/−^*, 62.2; p = 0.0062). Although the survival rate of *panx1b^−/−^* larvae compared to TL improved by day 6 (values in %: TL, 83; panx1b^−/−^, 87; p = 0.538; Fig. 2a), this phenomenon was unusual. The visual inspection of the gross anatomy of larvae after hatching on 3dpf, showed that *panx1b^−/−^* mutants developed a spectrum of morphological phenotypes from normal to malformed. Malformations included swimming bladder edema, ventricular hypertrophy, an irregularly shaped yolk sac, and abnormal spinal curvature in approximately 11±4% of mutants. Whether these malformations derived from abnormal development, or the microinjection process was not determined since larvae representing this spectrum of malformations could not survive beyond 5-6 dpf. **Fig. 2b** represents *panx1b^−/−^* and TL control larvae of the F4 generation at 6dpf. At this stage morphological malformations were not apparent. However, the head-to-tail size of 6dpf *panx1b^−/−^* larvae was significantly larger than the size of age matched TL controls (TL control = 3.83±0.27 mm; *panx1b^−/−^* = 4.00±0.17 mm; p < 0.0001; for TL, n = 60; *panx1b^−/−^*, n = 44; **Fig. 2c**). Adult *panx1b^−/−^* zebrafish were viable and fertile, with no apparent anatomical abnormalities. No gross morphologic changes to the eyes were observed. Furthermore, no significant reduction in the *panx1b* mRNA level was detected by real-time quantitative PCR (RT-qPCR) in 6dpf *panx1b^−/−^* larvae. This indicated that the 11 nt deletion in *panx1b^−/−^* RNA did not activate cytoplasmic RNA decay pathways like those activated in *panx1a^−/−^* (Safarian, Whyte-Fagundes et al. 2020). A compensatory regulation of the expression of the other three pannexin genes, *panx1a*, *panx2*, or *panx3*, was not detected (**Fig. 2d**). A prove of the loss-of Panx1b protein expression in zebrafish tissues was not achieved after several attempts to generate high-affinity polyclonal or monoclonal antibodies directed against both GST-*Panx1b* fusion proteins or synthetic peptides failed to generate a product fit for western blot analysis.

**Figure 2.**
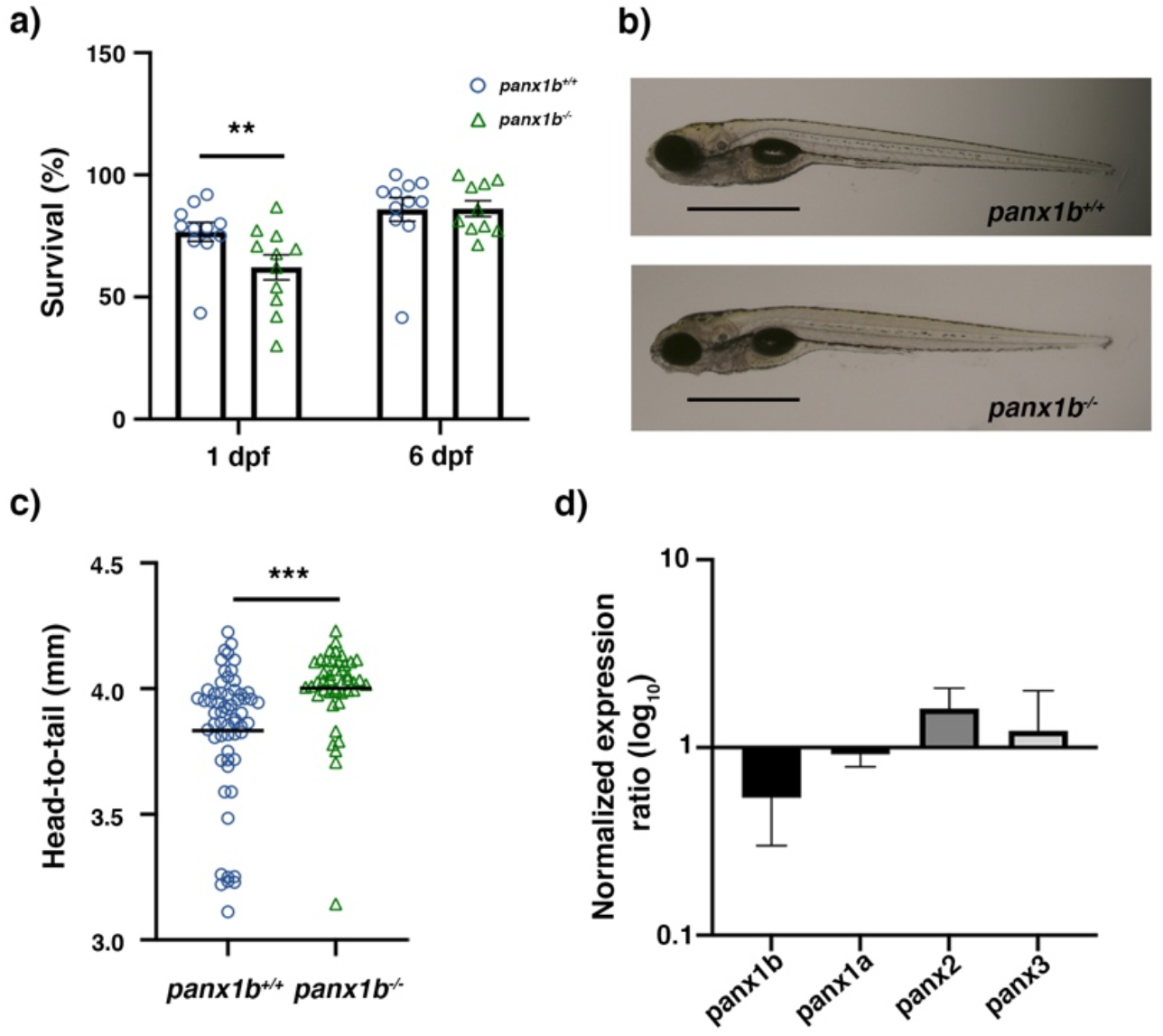
Primary characterization of panx1b^−/−^ larvae. Survival rate at 24 hours and six days post-fertilization (1dpf, 6dpf). b) Morphology of age-matched TL control and *panx1b^−/−^* larvae (at 6dpf) showing regular anatomical features. Scale bar = 100μm c) Head-to-tail size measurement of *panx1b*^−/−^ and control TLs (6dpf). d) RT-qPCR analysis of pannexin gene expression in 6dpf larvae. Significance: ** p<0.01 and ***p<0.001; nd: Not detected. Error bars = SEM.

### Transcriptome profiling of 6 dpf panx1b^−/−^ larvae

The transcriptomes of *panx1b^−/−^* and TL larvae were compared at 6 dpf following RNA-sequencing (NCBI Gene Expression Omnibus (GEO) database; at the time of submission RNA-seq data are scheduled for deposition at the GSE). A total of 58,616 RNA sequence tags representing the 25592 protein coding genes (Ensembl GRCz11 reference genome) were found (TL control, *panx1b^−/−^*; n=3). In *panx1b^−/−^* larva, 1213 RNAs were downregulated, and 764 RNAs were upregulated when the cut-off for the false discovery rate (FDR) was set to 0.05, and the significance of regulation was defined as p-value <0.001. Gene-specific expression information was retrieved from the Zebrafish Information Network (ZFIN) database to compare all regulated genes to those previously described as expressed in the central nervous system (CNS) and the visual system. Sixty-two upregulated genes and 92 downregulated genes were matched to expression in the central nervous system (**Fig. 3a**, left panel). Furthermore, 43 upregulated genes and 77 downregulated genes matched the genes expressed in the visual system (**Fig. 3a**, right panel).

**Figure 3.**
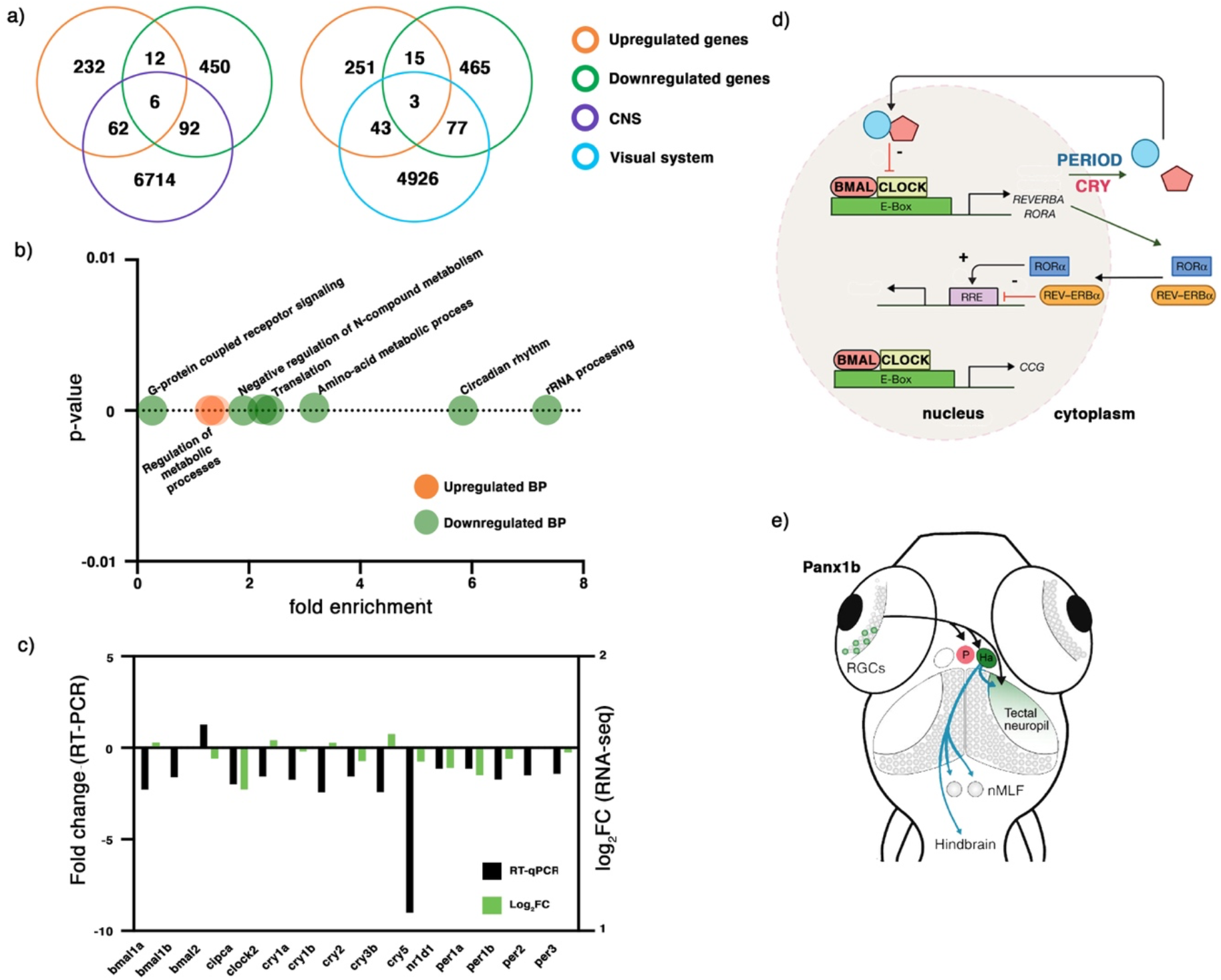
RNA-seq analysis of 6dpf panx1b^−/−^ larvae. **a)** Comparison of deregulated genes against two categories: central nervous system (CNS) and the visual system. **b)** GO annotation of RNA-seq data in terms of biological process (BP): the upregulated genes were enriched in the regulation of metabolic processes, whereas the downregulated genes were annotated for RNA processing, circadian rhythms, amino-acid metabolic process, translation, negative regulation of nitrogen (N) compounds metabolism, and G-protein coupled receptor signaling. **c)** Validation of RNA-seq data by RT-qPCR. The significant downregulation of *cry1, cry5, bmal1a, bmal1b*, *and clock2* genes in 6 dpf *panx1b^−/−^* larvae was confirmed by RT-qPCR in n = 3 independent replicates. **d)** Cartoon illustrating the network of circadian clock genes differently regulated in *panx1b^−/−^* larvae. Regulated genes in bold. **e)** Illustration of the connection of the inner retina with optic tectum, habenula (Ha), and pretectum (P) conferring light sensitivity and circadian rhythmicity, and descending control of locomotion. Abbreviation: nMLF, nucleus of the medial longitudinal fasciculus.

The AmiGO2 analysis (http://amigo.geneontology.org/amigo/landing) revealed enriched Gene Ontology (GO) terms and pathways (**Supplementary Tables 1 and 2**). The most significantly enriched biological processes (cut-off: FDR<0.05, p-value <0.001) represented the regulation of primary and macromolecules metabolic processes (**Fig. 3.b**, left panel). Downregulated genes represented several biological processes, including the circadian regulation of gene expression, RNA processing, and G-protein coupled receptors signaling pathways (**Fig. 3.b**, right plot).

The downregulation of circadian clock genes observed in the RNAseq data (cut-off: p-value<0.001; FDR 0.05) were also validated by RNA-qPCR (cut-off: p-value<0.05). Selected transcription factors driving the rhythmic 24-h expression patterns of core clock components were confirmed as downregulated in panx1b^−/−^ larvae (**Fig. 3.c**, **Table 1**). The modulation of the circadian clock pathway in the absence of Panx1b suggests a mechanism that could affect physiological functions such as sleep-wakefulness cycle, locomotor activity, or metabolic processes in 6 dpf old larvae by altering the circuitry which connects retina outputs with midbrain nuclei and the optic tectum **(Fig. 3d,e)**.

**Table 1.** RT-qPCR results for transcripts associated with the circadian clock pathway.

| Gene | RT-qPCR<br>Expression<br>(REST) | SEM (REST) | p-value (REST) | Result |
| --- | --- | --- | --- | --- |
| *bmal1a* | -2.277 | 0.050 | 0.005 | Down |
| *bmal1b* | -1.609 | 0.053 | 0.019 | Down |
| *bmal2* | 1.273 | 0.135 | 0.246 |  |
| *cipca* | -1.988 | 0.060 | 0.055 |  |
| *clock2* | -1.568 | 0.063 | 0.017 | Down |
| *cry1a* | -1.749 | 0.109 | 0.004 | Down |
| *cry1b* | -2.43 | 0.086 | 0.002 | Down |
| *cry2* | -1.569 | 0.073 | 0.058 |  |
| *cry3b* | -2.42 | 0.052 | 0.11 |  |
| *cry5* | -9.002 | 0.033 | 0.0012 | Down |
| *nr1d1* | -1.142 | 0.065 | 0.279 |  |
| *per1a* | -1.142 | 0.062 | 0.451 |  |
| *per1b* | -1.732 | 0.159 | 0.245 |  |
| *per2* | -1.503 | 0.082 | 0.154 |  |
| *per3* | -1.422 | 0.184 | 0.289 |  |

### Loss of Panx1b affected visually guided locomotor activities of larvae

Physiological, molecular, and behavioral studies have demonstrated robust circadian rhythmicity in the zebrafish based on the visual system (Hirayama, Kaneko et al. 2005). Since the transcriptome data analysis provided evidence that the lack of Panx1b caused changes in both the visual and the circadian clock systems, three types of visually guided locomotion-based tests, e.g., 1) locomotion in a stimulus-free environment, 2) visual-motor response (VMR) (Emran, Rihel et al. 2008), and optomotor response (OMR) (Bilotta 2000), were investigated.

Baseline locomotor performance of the *panx1b^−/−^* and TL larvae were compared under constant light (Light-ON) or dark (Light-OFF) conditions. Under the Light-ON condition, both TL controls and *panx1b^−/−^* swam longer distances and more rapidly than during the Light-OFF condition. Both genotypes displayed similar locomotor activity in Light-ON (**Fig. 4a**). The average of the distance traveled (t = 0.286, df = 52.295, p-value = 0.7758; for n=30 larvae) and the velocity (t = 0.686, df = 50.325, p-value = 0.496; for n=30 larvae) were comparable between both cohorts. Both groups showed a similar preference for swimming in the outer zone relative to the central “open space” area of the well (**Fig. 4a; well illustration**).

**Figure 4.**
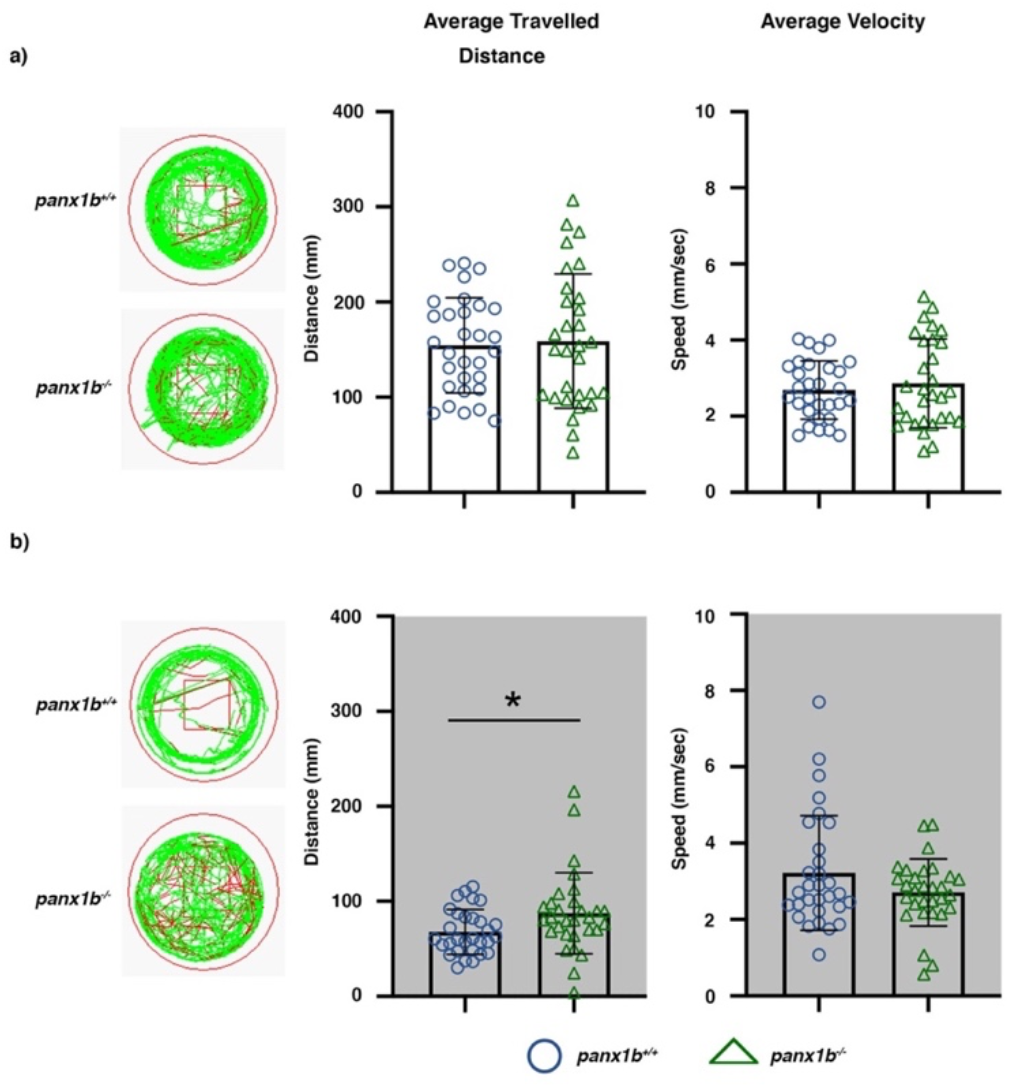
Locomotor activity in Light-ON and Light-OFF conditions. Locomotion was video tracked for 60 min in a) Light-ON and b) Light-OFF conditions. The wells on the left show examples of the locomotion patterns of *panx1b^−/−^* and *panx1b^+/+^* (TL control) larvae at 6 dpf. Medium (<20 mm/sec) or high-speed movements (>20 mm/sec) were visualized with green and red colors. Graphs demonstrate the averaged traveled distance (in mm) and the average velocity (mm/sec) of n = 60 larvae for each genotype. A Welch’s t-test was used for statistical comparison of groups. Significance: * = p-value<0.05. Error bars = SEM.

During the Light-OFF phase, *panx1b^−/−^* larvae were more active than TL controls. They swam significantly longer distances (t = 2.189, df = 45.194, p-value = 0.0338; n=30 larvae) although with similar velocity (t = −1.609, df = 46.751, p-value = 0.1142; n=30 larvae) as TL controls (**Fig. 4b**). No significant difference was observed in thigmotaxic behavior between both genotypes during the Light-OFF period; both groups stayed close to the walls of each well and avoided crossing the open space in the central zone (**Fig. 4b; well illustration**).

Next, the visual-motor response (VMR) assay was used to demonstrate the reaction of *panx1b^−/−^* larvae to sudden light changes. A principal component analysis (PCA) demonstrated that the two principal components PC1 and PC2 captured more than 65.5% and 29.5% of the data variance (**Fig. 5a**). The total activity duration (TAD; in red) had the most significant contribution to the observed variability in PC1 among seven variables tested (i.e., count and duration of freeze, bouts, and bursts of activity, plus total activity duration), (**Fig. 5b**).

**Figure 5.**
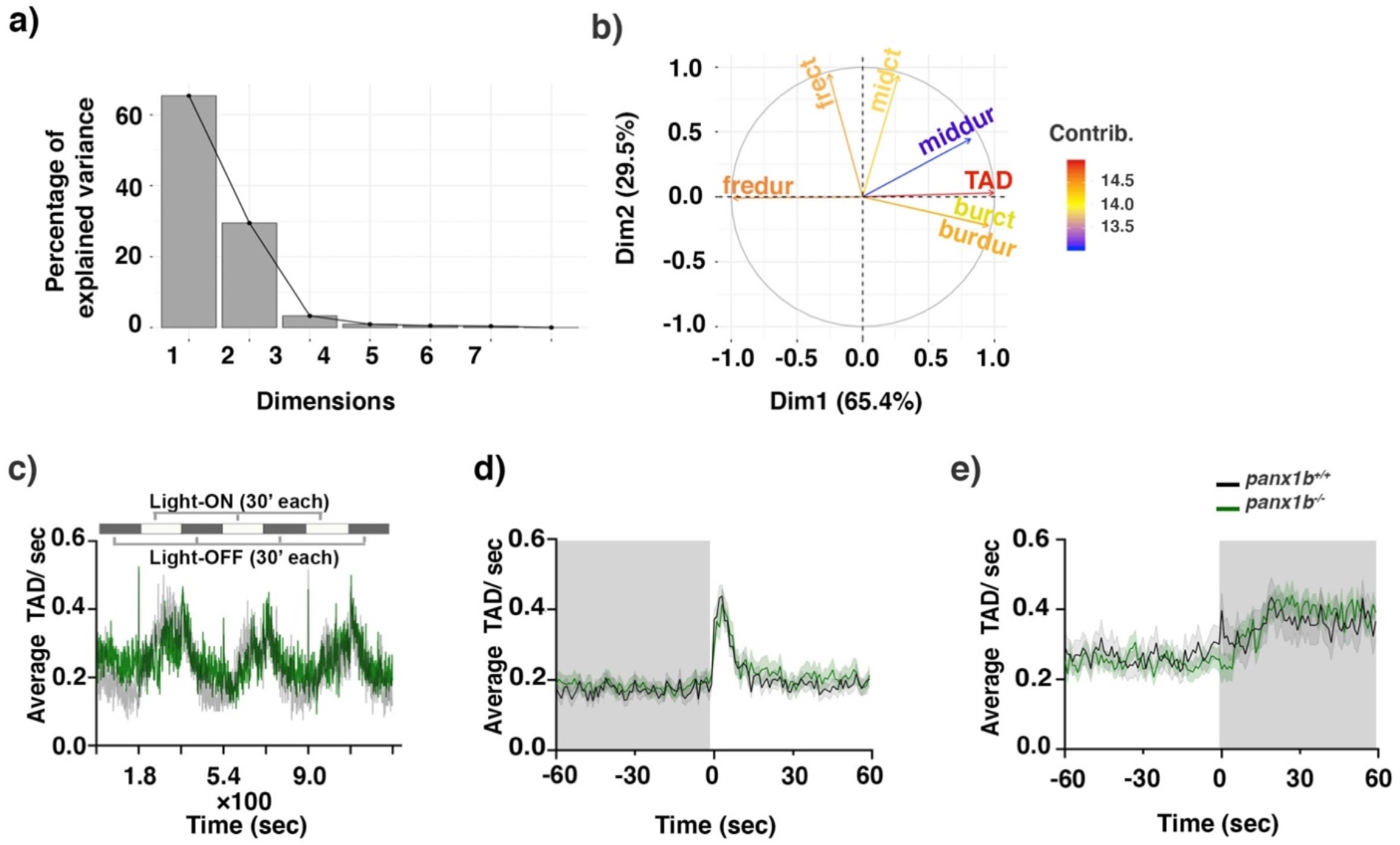
Visual-motor motor response (VMR) changes in panx1b^−/−^ larvae at 6dpf. a) Principal component analysis (PCA) transforming the VMR multidimensional data. The scree plot shows that Dimensions 1 and 2 capture more than 65.4% and 29.5% of the data variance. b) The variable correlation plot represented the coordinates of the variables in the first two dimensions. The variables were colored from blue to red as their contribution to PCs increases. Total activity duration (TAD; in color red) showed the highest contribution to the variability in Dimension 1 and was chosen to visualize differences between genotypes. c) The outline of the experimental paradigm is shown on top. The line graph shows the average TAD of *panx1b^−/−^* and *panx1b^+/+^* (TL) larvae from a representative test with n=24 larvae for each genotype. The activity was defined as the fraction of frames per second that a larva spent swimming (see methods for the detailed information). d, e) The results for average Light-ON and Light-OFF VMR are shown from 1 min before the light switch to 1 min after the light switch. The color ribbons surrounding the average activity line graph correspond to 1 S.E.M. The values are the average of the second trial from four independent tests with n=48 larvae for each genotype. The Light-ON and Light-OFF periods are indicated by white and black bars at the top of the panels. The “nparcomp” analysis was used for multiple comparisons and simultaneous confidence intervals.

The variable TAD was chosen to determine the VMR differences between both genotypes. The larval VMR differed from TL controls when the *panx1b* gene was ablated as indicated by the mean larval activity during 3 cycles of light onset (Light-ON) and light-offset (Light-OFF) (**Fig. 5c**). While the mutants’ motor activity during the Light-ON phases was markedly lower than TLs, they had comparable activity levels during the dark periods. A pairwise non-parametric t-test (nparcomp package; R) quantified larval activity during the following time periods: −60–0 sec (i.e., before light change), 0-1 sec (i.e., at the light change stimulus) and 1–60 sec (i.e., after light change). For the 1 min period before the Light-ON stimulus (−60-0 sec), there was a statistical difference in the activity between TL and *panx1b^−/−^* (p-value=0.001), in which the activity level of *panx1b^−/−^* before the light change was greater than that of TL larvae. At the dark-light transition (Light-ON; 0-1 sec), a synchronous increase in locomotor activity was observed in both TL and *panx1b^−/−^* groups (p-value= 0.083), which was followed by a rapid activity decline to a baseline level during the subsequent light period (in 1-60 sec, p-value<0.699; **Fig. 5d**).

At the light-dark transition, a synchronous and abrupt increase in motor activity was observed in TL control larvae. Following this trend, larval motor activity remained elevated for several minutes before slowly declining within 10 – 15 minutes, until swim activity fell below the baseline activity level in Light-ON conditions (**Fig. 5e**). Interestingly, *panx1b^−/−^* larvae responded to the dark differently compared to control TLs. For the 1 min period before the Light-OFF stimulus (−60-0 sec), *panx1b^−/−^* larvae activity was markedly lower than the activity of TLs (p-value<0.0001). Notably, the peak response to the abrupt loss of illumination was absent in mutants (0-1 sec; p-value<0.0001). Rather than sharply increasing motor activity, mutants responded to the Light-OFF by freezing for several seconds, indicated by periods of no activity. Then, activity gradually increased to levels comparable to TLs in the subsequent dark period. For the 1 min period after the Light-OFF stimulus, no significant differences between TL and the mutant’s activity were observed (1-60 sec; p-value=0.06) (**Fig. 5e**). The results showed that *panx1b^−/−^* larvae can detect and respond to light. However, their peak response to a dark pulse was abolished, which suggested that Panx1b in the retina functions in detecting light decrement and eliciting the corresponding swimming response in larval zebrafish.

Finally, the OMR test was used to assess the innate behavior of zebrafish, enabling them to follow visual motion in the surrounding environment to stabilize their location or course of movement (Neuhauss 2003, Portugues and Engert 2009). The optimal stimulus parameters for distinguishing mutants from TL control larvae were found by presenting moving gratings at three spatial frequencies (SF; 64, 128, and 256 pixels/cycle), and two velocities (72 and 144 deg/s), and two contrast levels of (10, 100 percent). **Fig. 6** shows the percentage of positive response (PPR) of 40 larvae after 90 s of exposure to grating motion towards alternating right and left stimuli. For all combinations of SFs, velocities, and contrasts, the percentage of *panx1b^−/−^* larvae responding to the stimulus was lower than that in the TL control group. A bias was detected when the direction of moving stripes was set towards the left side.

**Figure 6.**
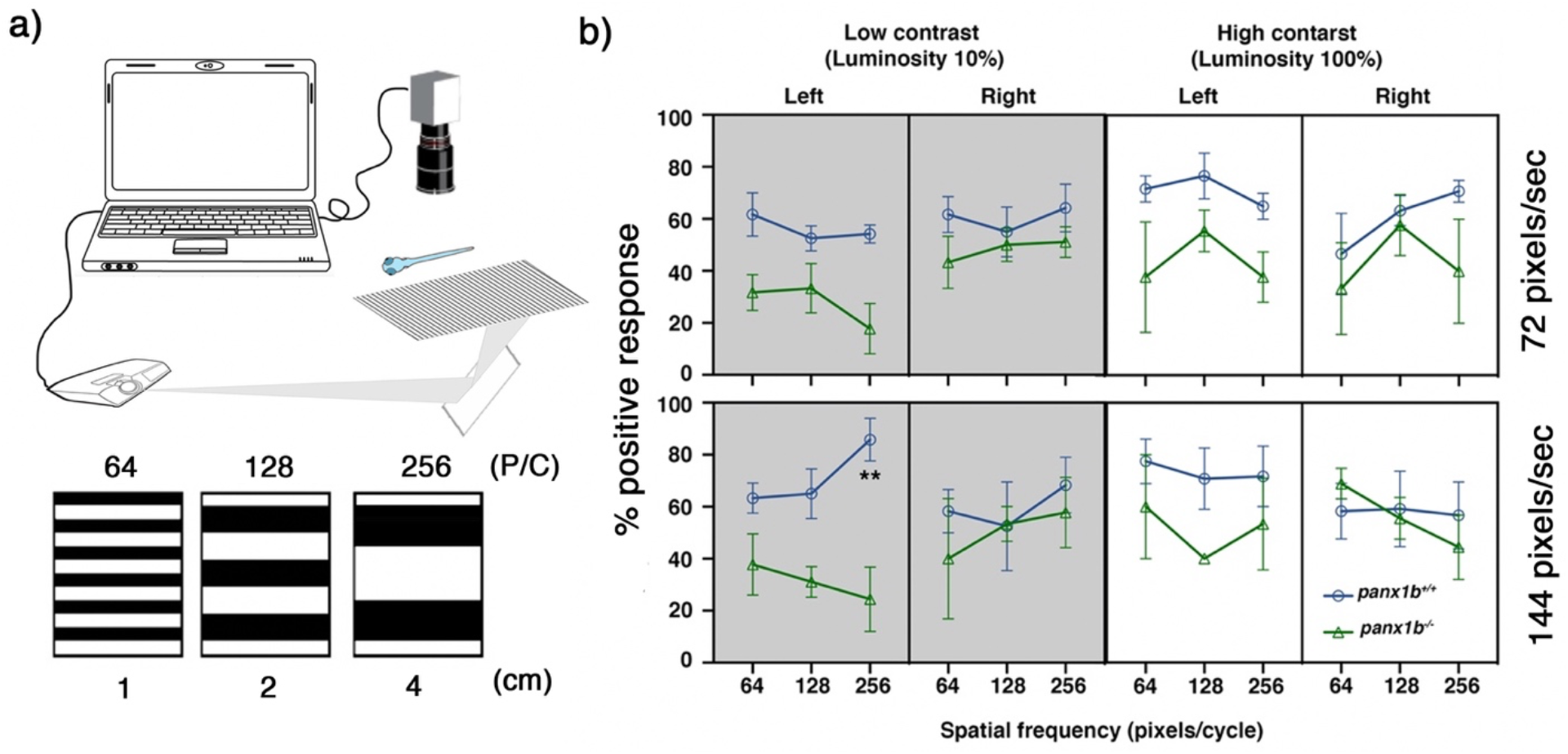
Optomotor response (OMR) declines in the absence of functional Panx1b. **a)** The OMR setup. Three spatial frequencies at two different speed rates and two luminosity levels were used to assess larval zebrafish’s visual acuity. P/C = pixel/cycle **b)** The percentage of positive response (% PPR) explains the average number of larvae that responded to a particular setting. The PPR values for each setting are the average of four tests ±SEM. In all tested settings, *panx1b^−/−^* were able to elicit an appropriate response, as observed in TL controls. However, their visual acuity was lower than controls when the direction of motion was towards the left. The difference between genotypes’ PPR for the left-ward motion became very significant at a spatial frequency of 256, with a speed of 144, and a low contrast setting (n=40). The “lm” and the model statistical “summary” functions were used to fit a regression model. The significant differences were calculated using the “TukeyHSD” function in R. Significance: p-value<0.05. Error bars = SEM.

At low contrast level (10%; left panels), *panx1b^−/−^* larvae were markedly distinct from TL controls when tested with left-ward moving stripes at a spatial frequency of 256 pixels/cycle and a speed of 144 pixels/s. The PPR of TL and *panx1b^−/−^* larvae were 85.8±16.4% and 24.4±21.4%, respectively (p-value=0.0012; n=40). For the low-spatial frequencies (i.e., 64 and 128 pixels/cycle), both genotypes swam most often in the direction of the moving stimulus when tested with a velocity of 72 or 144 (deg/s). No significant differences were observed within or between groups. However, when spatial frequencies were increased from 128 to 256 pixels/cycle, the average PPR was augmented at high velocities in TL controls. This result was in line with the previous reports on the larval zebrafish visual acuity peaking at the spatial frequency of 256 (pixels/cycle), and at a speed of 144 (deg/s) (Stiebel-Kalish, Reich et al. 2012). A similar increase in the larval response was observed in mutants only when larvae were exposed to the rightward moving grating. *Panx1b^−/−^* larvae showed a significant impairment for following the leftward moving stripes at the same combination of SF and velocity. The data suggested that the visual acuity of *panx1b^−/−^* larvae was compromised in low contrast conditions, which was in line with the aforementioned VMR result in the dark environment. Furthermore, the peak response deficiency, observed at SF of 256 pixels/cycle and speed of 144 pixels/s, for the leftward direction provided first evidence that Panx1b could be involved in a directional tuning impairment in *panx1b^−/−^* larvae. This advocates for a role of Panx1b in the processing of motion direction signals.

On the other hand, at high contrast level (100%), none of the tested combinations of SFs and velocities elicited a statistically significant difference between mutants’ and TLs’ OMRs (**Fig. 6, right panels**). The pairwise comparisons for each spatial frequency revealed no statistically significant differences within groups for either velocity. This outcome suggested that in a high contrast condition, when the environment was bright and the edges of objects were well defined, the visual acuity of *panx1b^−/−^* larvae was normal (p-values: see **Table 2)**.

**Table 2.** The multivariate comparisons of OMR between TL control and panx1b^−/−^ larvae.

| Contrast levels |  | Low (10%) |  |  | High (100%) |  |  |
| --- | --- | --- | --- | --- | --- | --- | --- |
| Speed<br>(pixels/sec) | Direction | Spatial Frequencies (pixels/cycles) |  |  |  |  |  |
|  |  | 64 | 128 | 256 | 64 | 128 | 256 |
| 72 | Leftward | 0.061 | 0.51 | 0.16 | 0.23 | 0.22 | 0.22 |
|  | Rightward | 0.52 | 0.77 | 0.44 | 0.89 | 0.8 | 0.33 |
| 144 | Leftward | 0.22 | 0.13 | 0.0012 | 0.63 | 0.26 | 0.66 |
|  | Rightward | 0.97 | 0.97 | 0.97 | 0.65 | 0.99 | 0.77 |

## Discussion

The localization of Panx1b in the inner retina and the ganglion cell layer offers opportunities to study pannexin functions in the retinal output pathway and in visual behaviors. Here, transcription activator-like effector nucleases (TALEN) introduced an 11-bp deletion in exon 4 of the *panx1b* gene causing a loss-of function mutation in the Panx1b protein. The strategy used is comparable to the conditional gene targeting of protein coding exons 3 and 4 in a Panx1 knockout mouse model (Dvoriantchikova, Ivanov et al. 2012, Prochnow, Abdulazim et al. 2012) and to our other *panx1a−/−* and *panx1b*^−/−^ zebrafish models (Cenedese, de Graaff et al. 2017, Safarian, Whyte-Fagundes et al. 2020). Like adult *panx1a^−/−^* the *panx1b*^−/−^ zebrafish were viable and fertile, with no apparent anatomical abnormalities or visible morphologic changes to the eyes interfering with the study design.

At 6dpf the visual system and visually guided locomotor activity of larvae is well established (Neuhauss 2003). This allowed to test three types of visually guided behaviors. The absence of Panx1b impaired both visually guided locomotor activities and the visual-motor response (VMR) of larvae in the dark differently to the changes when Panx1a is lost (Safarian, Whyte-Fagundes et al. 2020). The pattern of immediate freeze followed by partial recovery following the Light-OFF stimulus was distinct from the sustained reduction in locomotion seen in *panx1a^−/−^* larvae (Safarian, Whyte-Fagundes et al. 2020). Supporting evidence for a function in OFF-type retinal ganglion cells (RGCs) derives from patterned electroretinograms (PERG) and patch-clamp recordings of mice with targeted ablation of the Panx1 gene (Dvoriantchikova, Pronin et al. 2018). The response of OFF-type RGCs in mice and our data in the zebrafish let us conclude that the immediate response to sudden darkness could be associated with a similar disruption of the inner nuclear layer circuits, where Panx1b is localized (Kurtenbach, Prochnow et al. 2013).

The third behavioral test, the optomotor response (OMR) assay, detects changes to swimming posture testing the ganglion cells’ function (Portugues and Engert 2009, Naumann, Fitzgerald et al. 2016, Kist and Portugues 2019, Kramer, Wu et al. 2019). Without prior training and while larvae were freely moving, we showed robust OMR at 6 dpf in line with other studies presenting this response in larvae as early as 5 dpf (Portugues and Engert 2009, Portugues and Engert 2011). Here, we demonstrate that Panx1b affects the optic flow’s direction-selectivity, affecting the left-ward direction of motion at high spatiotemporal frequencies and low contrast conditions.

Presently, we can only speculate about how Panx1b affects the direction of motion. Future research must explain whether a similar right-ward motion detection exists, and whether Panx1a could play a role in it. In both zebrafish and in humans, motion detection is dominated by a luminance channel, which pools red and green cone signals (Orger and Baier 2005). However, RNA-seq data exclude the differential expression of red-or green-sensitive opsins, which differs to the upregulation of blue and UV-sensitive opsins and genes representing the phototransduction cascade found in *panx1a^−/−^* (Safarian, Whyte-Fagundes et al. 2020). While this observation does not conclusively rule out altered photoreceptor function, roles of Panx1b are more likely to affect the inner retina based on the localization in ganglion cells and end feet of Muller glia cell.

In the inner retina direction-selective ganglion cells (DSGCs) process and encode the presence, time, and the direction of motion (Naumann, Fitzgerald et al. 2016). Zebrafish RGCs send a diverse and highly regionalized time and spectral code to the brain (Zhou, Bear et al. 2020). The direction of motion is encoded by three direction-selective subtypes of ganglion cells tuned to upward, downward, and caudal-to-rostral motion (Abbas, Triplett et al. 2017). Furthermore, the direction tuning of multiple subtypes of DSGCs depends on the light level (Tikidji-Hamburyan, Reinhard et al. 2015). The direction selectivity of subpopulations of ON-OFF DSGCs in mammalian models requires electrical coupling by gap junctions (Vaney 1994, Trenholm, McLaughlin et al. 2013, Trenholm, McLaughlin et al. 2014), striking a balance between improving the detection of motion in dimly lit conditions against a cost to the accuracy of direction estimation (Yao, Cafaro et al. 2018). Since the OMR deficiency in the absence of Panx1b occurred only with the left-ward motion at low light conditions, it is intriguing to speculate whether Panx1b channels located in the inner plexiform layer play similar roles or synergize with connexins in broadening direction tuning at low light levels in specific DSGCs subtypes.

Astrocytes can modulate the activity of neurons *in vivo*, affecting directly information processing and suppressing motor output in the OMR test (Mu, Bennett et al. 2019). Astrocytes are also a main source of extracellular purines such as ATP and adenosine (Halassa, Fellin et al. 2009) (Bazargani and Attwell 2016). In rodents Panx1 is a major conduit of non-vesicular ATP release into the extracellular medium ((Suadicani, Iglesias et al. 2012, Beckel, Argall et al. 2014), and astrocytes release ATP in a circadian manner (Marpegan, Swanstrom et al. 2011). The localization of Panx1b in Muller glia end feet is close to the axonal output from ganglion cells and the retinal vasculature. In our opinion, a hypothetic model based on a photoperiodic ATP release involving both neurons and glia cells is attractive in explaining the observed behavioral phenotypes in the zebrafish, or the regulation of non-image forming processes such as sleep-wakefulness, which has been demonstrated in Panx1 mouse models (Kovalzon, Moiseenko et al. 2017, Shestopalov, Panchin et al. 2017). We propose a role of Panx1b in regulating clock-gene mediated heightened retinal sensitivity, which improves luminance detectability. This role fails to function properly in *panx1b^−/−^* larvae, leading to a diminution of the Light-OFF stimulus response.

To the best of our knowledge, we report a new twist to the expanding repertoire of Panx1 functions. The results presented indicate that lack of Panx1b compromises the final output of luminance and direction of motion detectors. Also, mutant larvae show a deficiency in following leftward direction of motion in low light conditions. The observed misalignment of circadian clock system seems to disrupt synchronization of behavioral rhythms with the circadian light-dark cycle in the absence of Panx1b. The principal findings in the zebrafish offer opportunities for studies on the Panx1b association with the circadian clock pathway in a diverse array of biological processes beyond the visual system, such as sleep/wakefulness, or metabolism.

## Materials and Methods

### Experimental Animals

All zebrafish (*Danio Rerio*) of strain Tupfel long fin (TL) were maintained in a recirculation system (Aquaneering Inc., San Diego, CA) at 28⁰C on a 14hr light/10hr dark cycle. All animal work was performed at York University’s zebrafish vivarium and in S2 biosafety laboratory in accordance with the Canadian Council for Animal Care guidelines after approval of the protocol by the Animal Care Committee (GZ: 2020-7-R3).

### Generating *panx1b* knockout zebrafish

Potential TALENs target sites were identified using Mojo Hand software (http://talendesign.org) (Neff, Argue et al. 2013). TALENs targeted exon 4 of *panx1b* (NM_001100030.2). The following criteria satisfied the TALEN design: TALENs target sites were 15-17 bases long with an initial 5’ T nucleotide to the TALE domain. The spacer length was restricted to 15-16 base pairs. Target sites with a unique restriction enzyme sequence located in the middle of the spacer sequence were selected to simplify screening for insertion-deletion (indel) mutations. The specificity of selected TALENs target sequences was determined using the BLAST interface built into the Mojo Hand software and by a NCBI BLAST search.

The TALEN constructs were synthesized in Dr. Stephen Ekker’s lab (Mayo Clinic Cancer Center, Rochester, MN). TALEN assemblies of the repeat-variable di-residues (RVD-containing repeats) were conducted using the Golden Gate approach (Cermak, Doyle et al. 2011). pT3TS-GoldyTALEN expression vectors (Bedell et al., 2012; A. C. Ma et al., 2013; A. C. H. Ma et al., 2016). Plasmids were linearized with the SacI restriction endonuclease (ThermoFisher Scientific, Canada) for 15 min at 37°C and used as templates for *in vitro* transcription. Capped cRNAs were synthesized from TALEN pairs mixed 1:1 using the mMESSAGE mMACHINE T3 Transcription kit (Life Technologies, Canada) and purified using the Oligotex mRNA Mini Kit (Qiagen Inc., Toronto, Canada). TALEN cRNAs were diluted in DNase/RNase-free water (Life Technologies) to the final concentration of 1 μg/μL and stored at −80 °C before microinjection.

One-cell stage zebrafish embryos were microinjected with TALEN cRNAs pair at a dosis of 25 pg/nl. Genomic DNA (gDNA) was extracted from injected embryos at 4dpf to examine the TALEN mutagenesis efficiency. Individual larvae were incubated in 100mM NaOH at 95℃ for 15 min. After cooling to room temperature, one-tenth of the 1 M Tris (pH8.0) volume was added to the extracts to neutralize the NaOH (Meeker et al., 2007). Finally, 1 volume TE buffer pH8.0 was added, and gDNAs were stored at −20℃. PCR was used as a screen to detect a small indel mutations followed by HindIII (for *panx1b*) restriction enzyme (RE) digests. PCR primers for genotyping: 5’-TTGGCGTCGGCAAACAGTGG-3’; reverse, 5’-GGCGACGAGGTGGTCGTTGG-3’. Indel mutations were confirmed by sequencing (Eurofins Genomics LLC, KY, USA) of gel-purified PCR products cloned into the pJet1.2 cloning vector (Life Technologies).

Adult mosaic zebrafish (F0) were anesthetized in pH-buffered 0.2 mg/ml ethyl 3-aminobenzoate methanesulfonate solution (MS-222, Sigma-Aldrich). The caudal fin (2 mm of the end) was removed using dissecting scissors (WPI Inc., FL, USA) and placed into 1.5 ml collecting tubes. The fin gDNA was isolated and screened for indel mutations as described (Bedell et al., 2012). Adult F0 zebrafish were out-crossed to wild-type (WT) TL zebrafish and their F1 offspring were analyzed by PCR and *HindIII* restriction digestions to verify germline transmission of mutations. Heterozygous *panx1b+/-* F1 mutants were in-crossed to establish homozygous F2 mutants *panx1b−/−*. All experiments described were performed with progenies of >F3 generations. Later generations were routinely tested for the identity of the genotype.

### Gene expression and transcriptome analysis

Total RNA was extracted from 6 dpf larvae (RNeasy Plus Mini Kit; Qiagen) and reversed transcribed (iScript Reverse Transcription Supermix; Bio-Rad Laboratories, Mississauga, Canada). The cDNA equivalent of 15ng total RNA was analyzed in triplicate by qRT-PCR (SsoAdvanced SybrGreen PCR mix; Bio-Rad). All experiments included a melt curve analysis of PCR amplicons generated in each reaction. Raw cycle threshold values (Ct-values) were exported from the CFX Manager Software (Bio-Rad, Canada), and the relative gene expression was calculated using the Relative Expression Software Tool (REST-2009) (Pfaffl, Horgan et al. 2002). All primer sequences (Integrated DNA Technologies, Toronto, Canada) are listed in the **Table 2**.

**Table 2.**
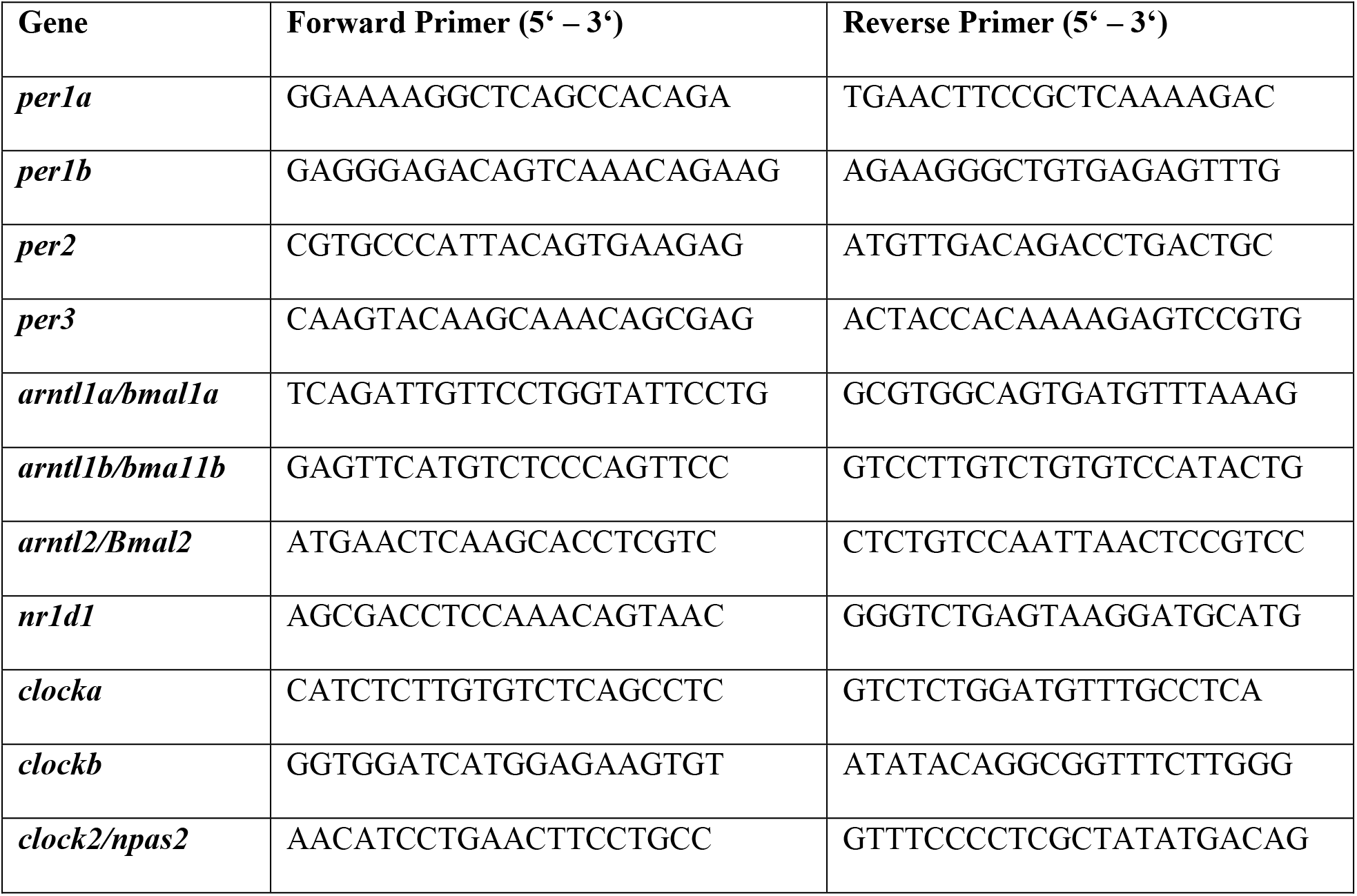
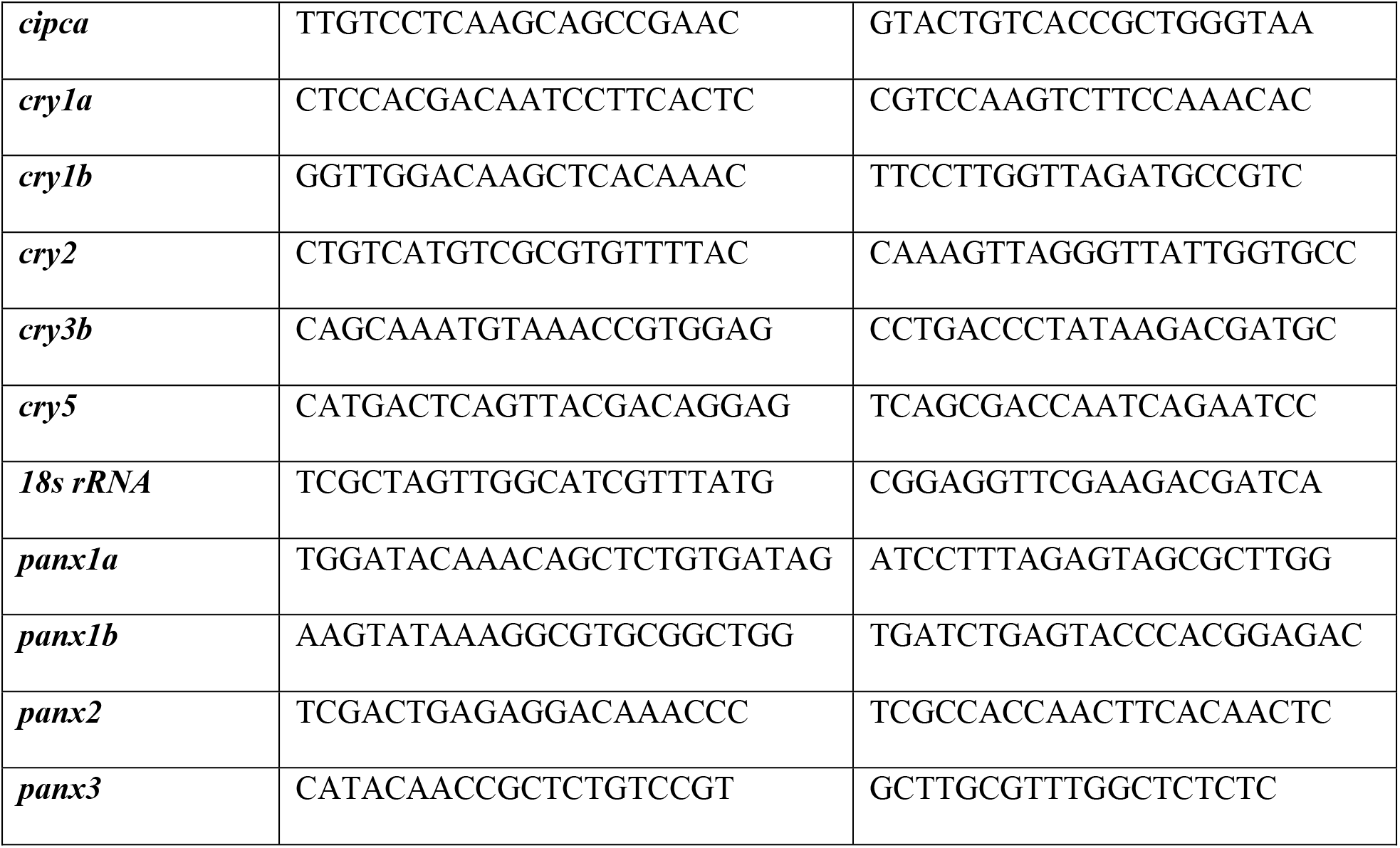
Primer sequences (RT-qPCR).

The transcriptomes of the zebrafish lines used in this research were analysed by RNA-seq (NGS-Facility, The Center for Applied Genomics, SickKids, Toronto, ON). The data derived from the sequencing of three independent pools of ≈30 age matched larvae (6dpf). Details of the bioinformatical analysis have been reported (Safarian, Whyte-Fagundes et al. 2020).

### Cell culture, Panx1b transfection, and imaging

Neuroblastoma 2a (Neuro2a) cells were cultivated in DMEM with 2mM glutamine, 1% nonessential amino acids (NEAA), 1% penicillin, and streptomycin (PS), and 10% fetal bovine serum at 37oC and 5% CO2 in a humid-controlled incubator. Cells were transfected with 200 ng total endotoxin-free plasmid DNA using the Effectene transfection protocol (Qiagen Inc., Valencia, CA, USA). Plasmid DNA encoded for C-terminally EGFP- or DsRed-monomer tagged Panx1b fusion proteins, containing the full-length protein-coding region of Panx1b (aa 1-422; Gene ID: 567417) or the truncated Panx1b aa 1-179 mutant. At 48h post transfected cells were fixed with 4% paraformaldehyde for 20 min at room temperature before washing with 1xPBS and mounted for imaging with ProLong^®^ Antifade mounting media (Thermo Fisher Inc., Mississauga, ON, Canada). The fixed samples were imaged using a Zeiss LSM 700 confocal microscope with a Plan-Apochromat 63x/NA 1.4 Oil DIC M27 objective using the ZEN 2010 program interface. Images were collected by line averaging (4x) at 2048⨯2048 pixel resolution.

### Panx1b antibody production

A polyclonal rabbit anti-drPanx1b antibody was generated against an affinity-purified GST-Panx1b fusion protein to target the carboxy-terminal amino acids 286-422 of Panx1b. The antibody was produced and affinity-purified by Davids Biotechnologie GmbH (code#ZOI-B2; Regensburg, Germany). The specificity in immunohistochemistry was confirmed using retina and brain tissues from adult TL, Panx1b, and Panx1a knock out fish. Control reactions without primary antibody complemented the tests. Despite extensive testing, the affinity-purified IgG antibody failed in all western blot conditions tested, like the chicken anti-Panx1b reported earlier (Kurtenbach, Prochnow et al. 2013).

### Immunohistochemistry

Adult zebrafish were humanely euthanized in MS-222 solution (0.02% w/v, Sigma-Aldrich). Tissues were removed and fixed in 4% paraformaldehyde (PFA) in 1xPBS overnight at 4oC, followed by cryoprotection in 30% sucrose in 1xPBS. After embedding in Tissue-Tek O.C.T compound 15μm sections were cut on a cryotome (Thermofisher). Samples were washed three times for 5 min with 1xPBS containing 0.1% Tween-20 (PBST) at RT. Unspecific binding sites were blocked with freshly prepared 5% normal goat serum (NGS, Sigma-Aldrich) in PBST for 1hr at RT℃. Following blocking, samples were incubated with primary antibody (1:50, affinity-purified rabbit anti-*panx1b* antibody, code#ZOI-B2, Davids Biotechnologie GmbH; 1:100, rabbit anti-GFAP, code AB5804, Sigma-Aldrich) overnight at 4℃. Subsequent washes with PBST were for one hour at 4℃. The Alexa 488 goat anti-rabbit secondary antibody (1:1500 in 1% NGS PBST, Life Technologies) was applied for one hr at RT℃. After 3 washed with PBST followed by one wash water, specimens were mounted on microscope slides using ProLong Antifade with DAPI (Thermofisher). Confocal images were collected using LSM-ZEN2 software (Zeiss LSM700 system; Carl Zeiss MicroImaging, Oberkochen, Germany) with Plan-Apochromat 20x/0.8 or Plan-Apochromat 63x/1.3 oil DIC M27 objectives. For comparison of wild type and knockout tissues settings were kept identical.

### Behavioral testing

The freely swimming behavior assay visual-motor response (VMR) was tested using larvae at 6dpf using a Zebrabox^®^ behavior recording system (ViewPoint Life Technology, Lyon, France; http://www.viewpoint.fr), and the Zebralab software (ViewPoint Life Technology, Lyon, France; http://www.viewpoint.fr). Tracking videos were recorded at 30 frames per second (fps) under infrared light illumination using a Point Grey Research Dragonfly2 DR2-HIBW. A lightbox provided infrared (for Light-OFF recording) or visible light (for Light-ON recording). All experiments were performed at 28°C. At least three independent experiments were performed for each assay type.

#### Freely swimming behavior assay

Larval swimming activity under constant Light-ON/OFF conditions was tested using 48-well plates. Larvae were adapted to the environment for 3hrs before recording begins. For tracking in light, larvae were adapted to ambient light (30% of final output intensity). For tracking in the dark, larvae were acclimatized to darkness (0% of final output intensity) for 3hrs. Next, locomotor behavior was tracked for 60 min. For analysis of locomotion, three thresholds were defined: slow (<2mm), medium (2-20mm), and fast (>20mm). The mean traveled distance (mm) and velocity (mm/sec) in two swim speeds *(medium and fast)* was used for statistical analysis.

#### The visual-motor response (VMR) assay

Larvae in 48-well plates were adapted to darkness (Light OFF) for 3hrs. After dark adaptation, baseline locomotion was recorded. Then, three trials of alternating light onset (Light ON) and light offset (Light OFF) periods with each period lasting 30 minutes, for a total of 180 min, followed. The Light-ON stimulus was set to 100%, and the Light-OFF stimulus was set at 0% of the final output intensity. In the Quantization^®^ mode of the Zebralab software threshold settings for activity detection in successive video frames were defined as no movement (<6), medium (6-20), activity burst (>20). Activity data were collected per second, with periods selected for analysis outlined in results.

#### The optomotor response assay

OMR assays were performed using a custom-built system, which follows the design described (Stih, Petrucco et al. 2019). Briefly, stimuli were presented from below using an ASUS P3B 800-Lumen LED portable projector (https://www.asus.com/ca-en/Projectors/P3B/). The fish movements were recorded using a USB 3.1 high-speed camera (XIMEA GmbH, Germany) equipped with a 35mm C Series Fixed Focal Length Lens (Edmund Optics Inc., USA). An 830 nm long-pass filter (Edmund Optics Inc., USA) was used to block infrared illumination coming from a source at the bottom of the test environment. The videos were recorded using XIMEA Windows Software Package (https://www.ximea.com/support/wiki/apis/XI-MEA_Windows_Software_Package). The visual stimuli were generated using the “Moving Grating” program, an online stimulus generator freely available at http://michael-bach.de/stim/. The Visual stimulus consisted of sequences of binary (i.e., “black and white” or “gray and white”) bars. The stimuli were generated with three spatial frequencies (64,128 and 256 pixels/cycles). Each spatial frequency was tested at two different speed rates (72 and 144 pixels/sec), and two contrast levels (i.e., 10 and 100% of the final contrast output on the program, resulting in “gray and white” or “black and white” bars, respectively). Stimuli were presented to larvae (n=10) for 90 sec, twice in each left or right direction. Larvae (6 dpf) were transferred to a 3 cm Petri dish (Thermo Scientific) and allowed to acclimatize for 5 min before video recording. To quantify the population response to optomotor stimuli, the Petri dish was subdivided into three zones: for each grating motion direction, the last section of the area was denoted as the “target zone.” The proportion of the larvae in the “target zone” at the end of each trial was counted in relation to the total number of fish and called the percentage of positive response (PPR). The difference in the PPR of the fish after a rightward or a leftward presentation of a stimulus is a measure of the strength of the optomotor response.

### Principal Component Analysis

VMR data represent seven variables: the counts and the duration of time larvae spend performing each of the three predefined swimming parameters (i.e., freeze, scoot, and burst) plus the total activity duration (TAD). In order to unambiguously evaluate the nature of behavioral responses to visual stimuli, the multiple components were disentangled by a principal component analysis (PCA) using the “FactoMineR” and “factoextra” packages. First, the PCA scores were Varimax rotated in accord with the standard PCA approach. The orthogonal configuration of varimax rotation guarantees the different components are uncorrelated, thus each represents an independent behavioral pattern. The resulting values termed “eigenvalues” signify the amount of variance explained by each of the components. To visualize and compare the size of the eigenvalues we used Scree plots. Only the first two components were retained for further interpretation since their eigenvalues were greater than 1 and they accounted for more than 90% of total variance in the data. Next, the variables contribution percentage to each of the two retained components were computed and plotted. The most informative variable, “TAD”, was compared between groups using nonparametric rank-based exact multiple contrast testing and simultaneous confidence intervals (MCTP/ SCI) (*nparcomp* package; http://www.R-project.org). Mean TAD values of the entire test period were plotted to show control and mutant larvae activity patterns. The activity data from 1 min before (−60 sec) to 1 min after (+60 sec) light changes were plotted separately to visualize the locomotor activity differences of the two genotypes upon after abrupt light switches. To visualize the effects of different drugs treatments on “frect” and “TAD” of larvae upon light switches, we used line charts. The baseline activity (from 1 min before light switches) were used to compare the amount of change occurring in any of the two mentioned activity features during the first 1 second after light switches.

### Statistics and reproducibility

Unless otherwise stated, all statistical analyses were performed in R software version 3.4.0 (http://www.r-project.org). A p-value<0.05 was considered statistically significant. The average values for the survival rate, body size, traveled distance, and velocity were compared between two groups using Student’s t-test (with equal variance) or Welch’s t-test (without equal variance) as indicated. Details of procedures and software packages used for the VMR analysis are described before (Safarian, Whyte-Fagundes et al. 2020). For the OMR data a multiple regression model was used to assess the statistical significance of the difference between groups’ PPR mean vectors. Two R packages, i.e., “tidyverse” and “car,” were used for the analysis. The normality and homogeneity of the data variance were checked by the “Shapiro-Wilk Test” and “Levene’s Test,” respectively. The “lm” and the model statistical “summary” functions were used to fit a regression model. Then, the “TukeyHSD” function was used to detect the significant differences. The sample sizes (n) reported corresponding to the number of larvae tested. The minimum number of independent experimental replicates was n ≥ 3.

## Acknowledgments

We thank Drs. Suzan El-Rass and Xiao-Yan Wen (Zebrafish Centre for Advanced Drug Discovery & Keenan Research Centre for Biomedical Science, Li Ka Shing Knowledge Institute, St. Michael’s Hospital, Toronto, Ontario, Canada M5B 1T8) for help with generating the Panx1b knockout line. Special thanks to Janet Fleites-Medina and Veronica Scavo for zebrafish husbandry.

## Author Contributions

Conceptualization, N.S., G.R.Z; formal analysis, N.S. G.R.Z; investigation, all authors; resources, G.R.Z.; writing—original draft preparation, N.S.; writing—review and editing, all authors; visualization, N.S., G.R.Z; supervision, G.R.Z.; project administration & funding acquisition, G.R.Z. All authors have read and agreed to the published version of the manuscript.

